# Named Entity Recognition and Linking: a Portuguese and Spanish Oncological Parallel Corpus

**DOI:** 10.1101/2021.09.16.460605

**Authors:** Vitor D.T Andrade, Pedro Ruas, Francisco M. Couto

## Abstract

Biomedical literature is the main mean of communication for researchers to share their findings. Since biomedical literature is composed of a large collection of text expressed in natural language, the usage of text mining tools to extract information from those texts automatically is of utmost importance. The problem is that the majority of the state-of-the-art tools were not developed to deal with other languages besides English, which in biomedical literature is even more critical since a significant part of health-related texts is written in the author’s native language. To address this issue, this work presents a deep learning NERL (Named Entity Recognition and Linking) system and a parallel corpus for the Spanish and Portuguese languages focused on the oncological domain. Both the system and the corpus are available at https://github.com/lasigeBioTM/ICERL_system-ICR_Corpus.

## Introduction

To extract knowledge from unstructured texts such as those in biomedical literature, it is necessary to apply text mining tools [14]. A text mining pipeline includes, among others, the tasks of Named Entity Recognition (NER) and Named Entity Linking (NEL), which corresponds to the recognition of entities mentioned in the text and the mapping of those entities to a given knowledge base (KB), respectively. Thus, text mining tools provide researchers and physicians a more effective way to obtain, integrate and interpret data from different sources [17] and to reduce the required time to process information [13, 16].

This work focuses on the NER and NEL tasks. Currently, there are different methods available for carrying them out, such as rule-based and deep learning methods. Rule-based methods include dictionary approaches, in which a given text is matched against a lexicon defined by experts, using string matching. Deep learning methods are based on artificial neural networks with multiple layers that learn a function responsible for mapping an input to an output. For example, in text mining, the input is usually word embeddings, which are vector representations of words [3]. Deep learning methods are the state-of-the-art for the NER and NEL tasks, but rule-based methods such as graph models can also be included in the state-of-the-art for the NEL task.

Flair is a framework that allows the combination of different pre-trained embeddings from language models to design a single model for the NER task [1]. Pre-trained language models are obtained by training a model on a large-sized corpus. These models can be further fine-tuned for specific text mining tasks involving different corpora than the training corpora [5]. BERT [4] is one of the most widely used pre-trained language models. It is organized in a multi-layer bidirectional transformer encoder which allows to draw dependencies between the input and output. This pre-trained language model was trained on different corpora, originating several variations, such as BioBERTpt, which was trained on Portuguese clinical and biomedical corpora and fine-tuned for the NER task [12].

An example of a graph model for the NEL task is the PPR-SSM [6]. PPR-SSM is based on the Personalized PageRank (PPR) algorithm. The PPR is a variation of the PageRank, an algorithm to rank the importance of webpages [10]. The PageRank algorithm involves the traversal of a graph, in which each node is a webpage, and the edges are links between pages. In PageRank, the jump is performed to a random page of the graph from a given page. In PPR, this jump is not random; it occurs to a chosen webpage. PPR has been used in NEL, and, in this case, the graph includes the candidates for the entities present in a given document [6]. The PPR-SSM uses the semantic similarity between the candidates for each entity and their information content (IC) to improve the results of the NEL task. In PPR-SSM, candidates with high semantic similarity with the other nodes and a high IC tend to have a better ranking in the disambiguation graph. The IC is influenced by the frequency of the presence of an entity in a corpus. A semantic similarity measurement (SSM) corresponds to the similarity value between entities using the relations defined in KBs.

Currently, the majority of text mining solutions are not designed to deal with other languages besides English [9] which is problematic, since only about 5% of world population has English as their native language^1^.

Several shared tasks have emerged to create state-of-the-art NER and NEL solutions for clinical text in recent years. Due to the significant amount of biomedical literature being published in Spanish and the number of Spanish speakers globally, this language has been the focus of several of these shared tasks. One example of these tasks is CANTEMIST [8], whose goal is to recognize entity mentions of tumor morphology in Spanish documents and to assign them to the respective “Clasificación Internacional de Enfermedades para Oncología” (CIE-O) codes. However, the number of shared tasks and text mining tools for Portuguese text is significantly lower when compared to Spanish.

Since there is a high lexical similarity between Portuguese and Spanish [2, as cited in [15]], this work intends to validate the possibility of using similar text mining tools for Portuguese and Spanish and by transferring annotations between the two languages. To do so, we developed the ICERL (Iberian Cancer-related Entity Recognition and Linking) system, a NERL (Named Entity Recognition and Linking) system that uses deep learning, and the parallel corpus ICR (Iberian Cancer-related) corpus. Both are focused on the oncology domain. Additionally, we present the evaluation of the NERL system’s performance on the ICR corpus.

## Methods

### ICERL system

As it is shown in Figure 1, the ICERL system is composed of two parallel pipelines; one is designed for Spanish text and the other for Portuguese. The Spanish pipeline consists of the improvement of the pipeline that we initially designed for the CANTEMIST shared task [11] with a difference in the NEL task: instead of returning the best candidate from CIE-O, the system returns a list of candidates from CIE-O and DeCS.

**Figure 1.**
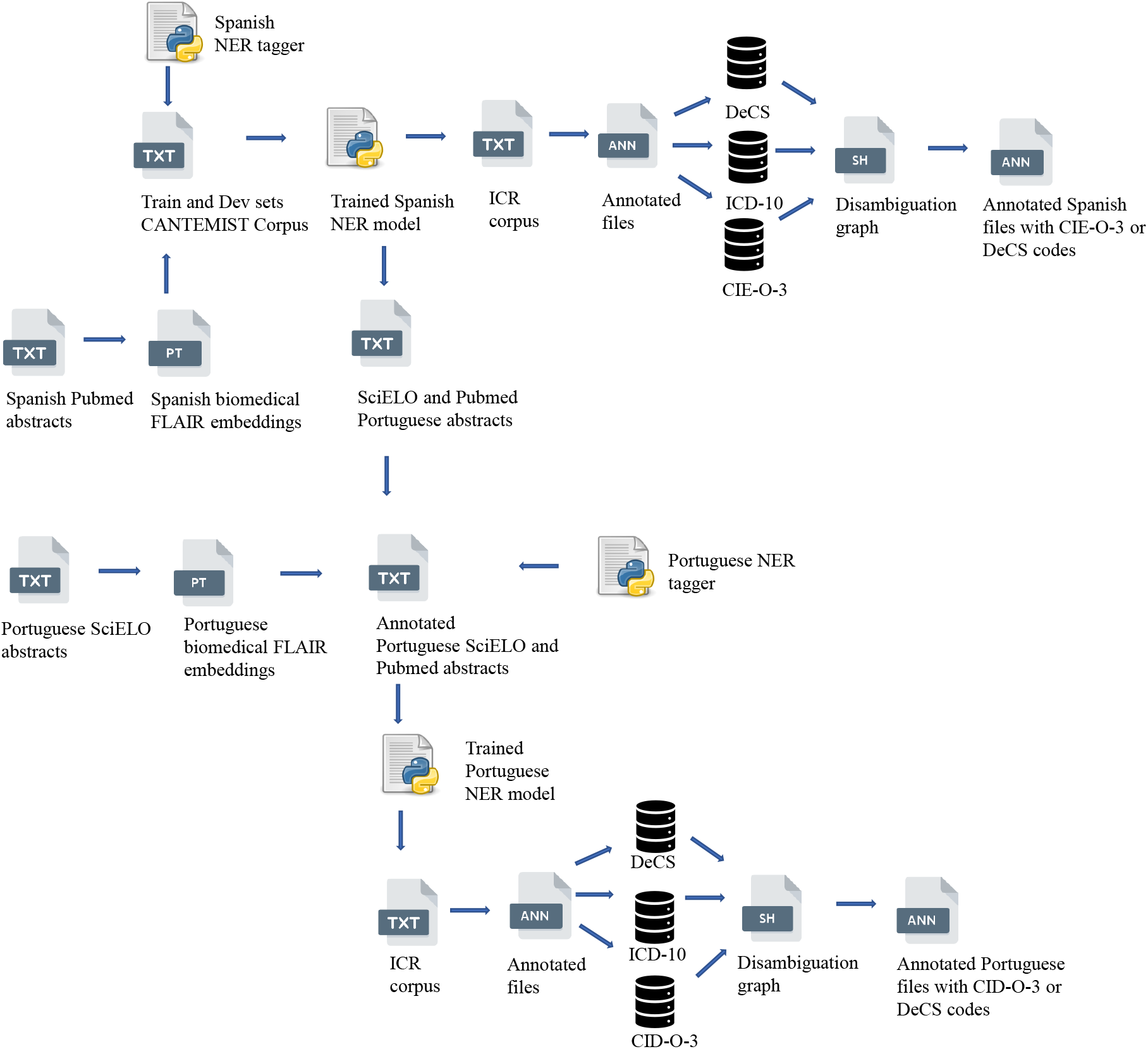
ICERL system.

The Portuguese pipeline is composed of similar embeddings to the ones used in the Spanish pipeline and the BioBERTpt model. In addition to making the Spanish annotations, the Spanish pipeline is also applied to the Portuguese training files. The resulting annotations and the Portuguese embeddings are used to train the Portuguese pipeline that is applied on the texts written in Portuguese.

### Spanish pipeline

This pipeline used the Flair framework for the training of biomedical embeddings and to build the NER model. For the embeddings, we used Flair embeddings (*es-forward* and *es-backward*) trained on Spanish Wikipedia, Spanish FastText embeddings, and Flair embeddings trained in 32,500 Spanish PubMed^2^ articles with 35,352,727 tokens. The Flair embeddings were trained with the following parameters: hidden size = 1024; nlayers=1; dropout = 0.1; sequence length = 250; mini-batch size = 32; max epochs = 71 epochs for the backward embeddings and 99 epochs for the forward embeddings and patience = 25. The resulting embeddings were integrated into a NER tagger that was further trained on the CANTEMIST corpus training and development sets [8] with the following training parameters: hidden size=256; learning rate = 0.1; mini-batch size = 32; max epochs = 55; patience = 3. The trained model was then applied to the CANTEMIST corpus test set to create the annotation files with the recognized entities.

Considering the entities present in the annotation files, the next step was to link them to the respective concepts in the CIE-O vocabulary (NEL task). Therefore, it was necessary to retrieve vocabulary candidates from CIE-O, which the CANTEMIST’s organization had provided, and also from DeCS and ICD-10. A disambiguation graph was constructed with these candidates, and through the PPR-SSM algorithm, the candidates were sorted, and the best candidate was linked to the entity mentioned in the text. Even though the entities could only be linked to CIE-O concepts, the role of the ICD and DeCS candidates was to create a complete disambiguation graph with more semantic information about the entities, which was expected to improve the precision of the disambiguation [11].

After replicating this pipeline, the next step was to improve it. For the NER task, we retrained the Flair embeddings up to 200 epochs, added Spanish BytePair and Character embeddings to the NER tagger, and retrained up to 200 epochs. For the NEL task, we fixed code errors in the string matching process, such as uppercase and lowercase letters, space between words in the entity string, and used the complete list of CIE-O codes and in CIE-O codes that were only available in the training and development of the CANTEMIST corpus. These modifications on the NEL task were inspired by SINAI’s approach, one of the teams that also participated in CANTEMIST [7].

### Adaptation to Portuguese

For the Portuguese pipeline, we developed three NER models:

- **”cantemistpt”:** it includes Flair embeddings (pt-forward and pt-backward) trained on Portuguese Wikipedia, Portuguese FastText embeddings, and SciELO Flair embeddings, Portuguese Bytepair embeddings, and Character embeddings.
- **”cantemistpt + biobertpt”:** it includes Flair embeddings (pt-forward and pt-backward) trained on Portuguese Wikipedia, Portuguese FastText embeddings, and SciELO Flair embeddings, Portuguese Bytepair embeddings, Character embeddings, and BioBERTpt embeddings.
- **”biobertpt”:** it includes Portuguese FastText embeddings and BioBERTpt embeddings.

The SciELO ^3^ Flair embeddings were trained in 500 articles with the following parameters: hidden size = 1024; nlayers=1; dropuout = 0.1; sequence length = 250; mini batch size = 32; max epochs = 118 epochs for the backward embeddings and 110 epochs for the forward embeddings and patience = 25. The three models were trained in SciELO’s articles related with oncological domain in Portuguese with hidden size=256; learning rate = 0.1; mini batch size = 32 (except the cantemist + biobertpt which had a mini batch size = 16); max epochs = 150; patience = 3.

### ICR Corpus

The parallel corpus is composed of 1555 articles in Portuguese and Spanish which were retrieved from SciELO. The following filter was used to obtain the articles: *((*) AND (oncology)) OR (cancer))* Furthermore, we ensured that only articles with abstracts available for both languages were included in the corpus.

In addition to the parallel corpus, other documents for the Portuguese pipeline were retrieved. The training of embeddings uses 500 abstracts retrieved from SciELO without any specific filter since these articles are intended to include several domains. For the training of the NER tagger, 974 abstracts were retrieved from SciELO and 41 articles from PubMed that were retrieved through the query *Case Reports[Publication Type] AND POR[LA] AND Cancer[Filter]*. Since this number of abstracts is small for the training of the parallel corpus, we resorted to nlpaug ^4^, a python library for data augmentation in order to increase the number of training documents. With this tool, we replace words with the respective synonyms from WordNet. For example, the word “cancer” is replaced by “câncro”. Thus, two new documents for each original document were created, thus making a total of 3045 documents for training. These documents were then annotated using the pipeline designed for the Spanish texts. These annotations were the same ones used for CANTEMIST; that is, each token from the training files was tagged with the label “B-MOR NEO” if it corresponded to the beginning of an annotation, the label “I-MOR NEO” if it corresponded to the inside of an annotation and the label “O” if the token was not part of any annotation. We then corrected the annotations manually. These corrections refer to errors in the tokenization and labeling processes. For example, in some instances, the entity “tumor” has split into two tokens(”tum” and “r”), and entities such as “cancer de boca” were labeled:

- ”cancer”: “B-MOR NEO”
- ”de”: “O”
- ”boca”: “O”

when it should be labeled:

- ”cancer”: “B-MOR NEO”
- ”de”: “I-MOR NEO”
- ”boca”: “I-MOR NEO”

### Evaluation

The ICERL system’s evaluation consists of comparing the results of the Portuguese pipeline with the results of the Spanish pipeline. The reason for this method of evaluation is because it is possible to assess the performance of the Spanish pipeline on the CANTEMIST corpus, and the results obtained are close to the state-of-the-art [8].

The results of the Portuguese pipeline correspond to a list of candidates from DeCS and CID and, the results of the Spanish pipeline to candidates from DeCS and CIE.. The usage of DeCS and a list of candidates instead of the best candidate served to not restrict the evaluation method to just one candidate and one vocabulary. For example, the entity “Neoplasia benigna” is in the DeCS vocabulary with the id “D009369” and in CID with the id “8000/0”. Thus, for each entity, this evaluation method considers a true positive if at least one of the candidates on the list in Portuguese is on the list of candidates in Spanish; a false positive if the Portuguese pipeline does not find an entity which was found by the Spanish pipeline or none of the candidates of the Portuguese entity is on the candidates of the Spanish entity; a false negative if the Spanish pipeline does not find an entity and the Portuguese pipeline does. With these instances, the three metrics to evaluate the ICERL system are calculated: precision, recall, and F1-score.

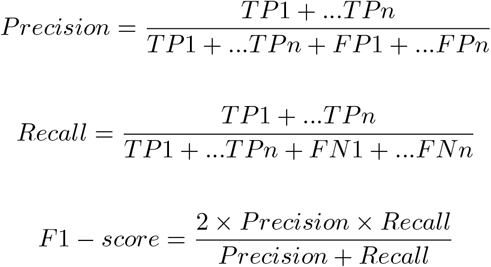

### True positive (TP) example

**ES sentence:** De los 52.912 casos, 83,4% eran mujeres y 96,9% era de carcinomas diferenciados.

**PT sentence:** Dos 52.912 casos, 83,4% eram femininos e 96,9% eram carcinomas diferenciados.

**ES entity:** carcinomas

**ES candidates:** [**2320**, 8270/3, **9081/3**, 8271/0, **8160/3, 8010/9, 8010/3**, 8140/3, **8934/3, 8337/3, 8231/3**]

**PT entity:** carcinomas

**PT candidates:** [**2320, 9081/3, 8160/3, 8010/9, 8010/3, 8934/3**, 8042/3, **8231/3**, 8102/3, 8300/3, **8337/3**]

### False positive (FP) example

**ES sentence:** Los resultados son consistentes con la epidemiología del cáncer de tiroides, con predominio del sexo femenino y carcinomas diferenciados.

**PT sentence:** Os achados são consistentes com a epidemiologia do câncer de tireoide, com predominância do sexo feminino e do carcinoma diferenciado.

**ES entity:** Not found

**ES candidates:** Not found

**PT entity:** carcinoma diferenciado

**PT candidates:** [2320, 8082/3, 8022/3, 8805/3, 8145/3, 8530/3, 8246/3, 8020/3, 2331, 38036, 9372/3, 2330, 31596, 31595, 9243/3, 2335, 2340, 34685, 8331/3, 31587, 2329]

### False negative (FN) example

**ES sentence:** Describir el perfil clínico y epidemiológico de los casos de cáncer de tiroides en Brasil

**PT sentence:** Descrever o perfil clínico-epidemiológico de casos hospitalares de câncer primário de tireoide no Brasil.

**ES entity:** cáncer

**ES candidates:** [8000/3, 9562]

**PT entity:** Not found

**PT candidates:** Not found

## Results and discussion

The modifications that were made in the Spanish pipeline [11] resulted in an improvement of the F1-score of NER task from 0.741 to 0.754, and from 0.061 to 0.665 on the NEL task. Even though that the improvements in the NER task were small, and the performance in the two tasks did not reach the state-of-the-art (F1-score of 0.87 on the NER task and 0.825 on the NEL task) [8], the improvements on the NEL task should be highlighted.

The application of the ICERL system on the ICR corpus resulted in 3,399 annotations for the Spanish language, of which 216 correspond to unique annotations and 3,287 in Portuguese, with 171 being unique annotations. The entity “cancer” was the most frequent annotation for both languages.

The Table 1 presents the results of the three models for the Portuguese language and the baseline. This baseline corresponds to the application of the Spanish pipeline to the texts in Portuguese. Even though the precision achieved by the baseline is high, the recall reveals that some of the entities that the system found in Spanish were not found in Portuguese, which means that it was possible to develop an approach with a better performance. To do so, the three models were developed. The results obtained by the three models show that the BioBERTpt and the adaption of the Spanish pipeline are both valid solutions for the recognition and linking of entities in our parallel corpus.

**Table 1.** Results of baseline and the three models for Portuguese text. P refers to Precision, R to Recall, and F1 to F1-score.

|  | **P** | **R** | **F1** |
| --- | --- | --- | --- |
| baseline | 0.887 | 0.171 | 0.287 |
| cantemistpt | 0.874 | 0.842 | **0.858** |
| cantemistpt + biobertpt | 0.873 | **0.844** | **0.858** |
| biobertpt | **0.898** | 0.821 | **0.858** |

The improvement in the recall and F1-score of these models when compared to the baseline, can be explained by the decrease of the FN instances, in other words, the number of times in which the system does not find an entity in Portuguese that it had found in Spanish significantly decreases. In addition to the differences regarding the embeddings between each model and the baseline, this improvement is also due to the manual correction of the Portuguese training files annotations. The final version of the ICERL system uses the “cantemist+biobertpt” for the Portuguese pipeline since this model is the one with the best recall and therefore it is the one with the smallest number of FN instances.

The NLPStatTest, a toolkit to compare the performance of two NLP systems, was used for the baseline and the final version of the ICERL system. The F1-score in each document was the score used to compare the two systems. As it is described in Table 2 the performance of the ICERL system is 58% higher than the baseline.

**Table 2.** Statistics of the baseline and ICERL system.

| **Score** | **Mean** | **Median** | **Std. Dev.** | **Minimum** | **Maximum** |
| --- | --- | --- | --- | --- | --- |
| ICERL system | 0.708 | 0.703 | 0.121 | 0.369 | 0.954 |
| Baseline | 0.121 | 0.089 | 0.118 | 0.000 | 0.578 |

To further assess the performance of the ICERL system, an error analysis was conducted in one hundred and eighty-three annotations. The entities that were not recognized by the Portuguese and Spanish pipelines correspond to 60.6% and 30.3% of the total errors. The other 10.1%, corresponds to the comparisons between different entities. For example, in the document “S0034-70942013000200006”, the Spanish annotation is “tumores epidermoides”, and the Portuguese is “tumores”. None of the candidates of the Portuguese annotation is in the list of candidates of the Spanish annotation, therefore the system considers this example as FP. The reasons for these three types of errors are due to the fact that different files were used to train the two pipelines, and the Portuguese training annotations were manually corrected.

The results obtained in Table 1 show that the ICERL system has a similar performance in both languages. Furthermore, the fact that there are no errors in which the same entities for Portuguese and Spanish do not have the same candidates reinforces this claim.

## Conclusions

This work culminated in two contributions, the ICR corpus, a Spanish-Portuguese cancer-related corpus, and the ICERL system, a NERL system for the oncological domain. These two resources are available at https://github.com/lasigeBioTM/ICERL_system-ICR_Corpus. The importance of these contributions lies in the fact that, as far as we know, there are no similar tools for Portuguese and Spanish there are cancer-related.

The similarity between the annotations statistics of the two languages and a 0.858 F1-Score achieved by the evaluation method confirms the initial hypothesis; it is possible to use similar text mining tools for Portuguese and Spanish and to transfer annotations between the two languages maintaining comparable performance.

For future work, we intend to decrease the annotations differences between the two languages. One of the ways to do so is to use a parallel corpus to train the ICERL system. This is because, despite manually correcting training annotations for the Portuguese pipeline, the amount of annotations does not allow all errors to be corrected. Moreover, the corrected training annotations for the Portuguese pipeline do not always follow the same annotation criteria used in the training files of the Spanish pipeline. The Spanish pipeline training files came from the shared task CANTEMIST and were designed only for entities present in the CIE-O vocabulary and not for DeCS. So, for example, the entity “neoplasia de la mama” is not present in the vocabulary CIE-O, but it can be found in the DeCS vocabulary.

The main problem with our approach is the time needed to train the models and the embeddings. For example, the model with a longer training time, the “cantemistpt + biobertpt” took three to four weeks to train. However, there was no significant improvement between training this model to 29 and 150 epochs, so in the future, we intend to find a number of epochs that optimizes the time spent for the training of each model.

In the future, it would also be interesting to extend the ICERL system and the ICR corpus to other Romance languages such as Catalan, Galician, Italian and French since there is a high lexical similarity between them and Portuguese and Spanish. In this case, the ICR corpus would have to be constituted by parallel corpora that included English in addition to these languages. The evaluation method of the ICERL system would be the comparison of the performance of these languages with a performance of a state-of-art tool on the English texts. Since there has been a greater focus on text mining tools for the English language, the performance obtained on texts in English will be higher than for other languages. Therefore, the results obtained on the texts in English will be considered as the gold standard.

## Footnotes

1 https://web.archive.org/web/20190312060544/https://www.ethnologue.com/statistics/size

2 https://pubmed.ncbi.nlm.nih.gov/

3 https://scielo.org/

4 https://github.com/makcedward/nlpaug

## References

1. A. Akbik, T. Bergmann, D. Blythe, K. Rasul, S. Schweter, and R. Vollgraf. Flair: An easy-to-use framework for state-of-the-art nlp. In Proceedings of the 2019 Conference of the North American Chapter of the Association for Computational Linguistics (Demonstrations), pages 54–59, 2019.

2. S. Castro, J. Bonanata, and A. Rosá. A high coverage method for automatic false friends detection for spanish and portuguese. In Proceedings of the Fifth Workshop on NLP for Similar Languages, Varieties and Dialects (VarDial 2018), pages 29–36, 2018.

3. F. M. Couto and A. Lamurias. Mer: a shell script and annotation server for minimal named entity recognition and linking. Journal of Cheminformatics, 10(1):58, 2018.

4. J. Devlin, M.-W. Chang, K. Lee, and K. Toutanova. Bert: Pre-training of deep bidirectional transformers for language understanding. arXiv preprint 1810.04805, 2018.

5. S. Edunov, A. Baevski, and M. Auli. Pre-trained language model representations for language generation. arXiv preprint 1903.09722, 2019.

6. A. Lamurias, P. Ruas, and F. M. Couto. PPR-SSM: Personalized PageRank and semantic similarity measures for entity linking. BMC Bioinformatics, 20(1):1–12, 2019.

7. P. López-Úbedaa, M. Diaz-Galianoa, M. Martin-Valdiviaa, and L. A. Urenã-Lópeza. Extracting neoplasms morphology mentions in spanish clinical cases through word embeddings. Proceedings of IberLEF, 2020.

8. A. Miranda-Escalada, E. Farré, and M. Krallinger. Named entity recognition, concept normalization and clinical coding: Overview of the cantemist track for cancer text mining in spanish, corpus, guidelines, methods and results. 2020.

9. A. Névéol, H. Dalianis, S. Velupillai, G. Savova, and P. Zweigenbaum. Clinical natural language processing in languages other than english: opportunities and challenges. Journal of biomedical semantics, 9(1):12, 2018.

10. L. Page, S. Brin, R. Motwani, and T. Winograd. The pagerank citation ranking: Bringing order to the web. Technical report, Stanford InfoLab, 1999.

11. P. Ruas, A. Neves, V. Andrade, and F. Couto. Lasigebiotm at cantemist: Named entity recognition and normalization of tumour morphology entities and clinical coding of spanish health-related documents. In Proceedings of the Iberian Languages Evaluation Forum (IberLEF 2020), 2020.

12. E. T. R. Schneider, J. V. A. de Souza, J. Knafou, L. E. S. e Oliveira, J. Copara, Y. B. Gumiel, L. F. A. de Oliveira, E. C. Paraiso, D. Teodoro, and C. M. C. M. Barra. Biobertpt-a portuguese neural language model for clinical named entity recognition. In Proceedings of the 3rd Clinical Natural Language Processing Workshop, pages 65–72, 2020.

13. C. Simon, K. Davidsen, C. Hansen, E. Seymour, M. B. Barnkob, and L. R. Olsen. Bioreader: a text mining tool for performing classification of biomedical literature. Bmc Bioinformatics, 19(13):57, 2019.

14. A.-H. Tan et al. Text mining: The state of the art and the challenges. In Proceedings of the PAKDD 1999 Workshop on Knowledge Disocovery from Advanced Databases, volume 8, pages 65–70. sn, 1999.

15. J. L. Ulsh. From spanish to portuguese. Foreign Service Institute, Department of State, 1971.

16. C.-H. Wei, H.-Y. Kao, and Z. Lu. Pubtator: a web-based text mining tool for assisting biocuration. Nucleic acids research, 41(W1):W518–W522, 2013.

17. F. Zhu, P. Patumcharoenpol, C. Zhang, Y. Yang, J. Chan, A. Meechai, W. Vongsangnak, and B. Shen. Biomedical text mining and its applications in cancer research. Journal of biomedical informatics, 46(2):200–211, 2013.

